# GSTZ1 sensitizes hepatocellular carcinoma cells to sorafenib-induced ferroptosis via inhibition of NRF2/GPX4 axis

**DOI:** 10.1101/2020.12.14.422655

**Authors:** Qiujie Wang, Bin Cheng, Qiang Xue, Qingzhu Gao, Ailong Huang, Kai Wang, Ni Tang

## Abstract

Increasing evidence supports that ferroptosis plays an important role in tumor growth inhibition. Sorafenib, originally identified as an inhibitor of multiple oncogenic kinases, has been shown to induce ferroptosis in hepatocellular carcinoma (HCC). However, some hepatoma cell lines are less sensitive to sorafenib-induced ferroptotic cell death. Glutathione S-transferase zeta 1 (GSTZ1), an enzyme in the catabolism of phenylalanine, has been found to negatively regulate the master regulator of cellular redox homeostasis nuclear factor erythroid 2-related factor 2 (NRF2). This study aimed to investigate the role of GSTZ1 in sorafenib-induced ferroptosis in HCC cell lines and determine the involved molecular mechanisms. Mechanistically, GSTZ1 depletion enhanced the activation of the NRF2 pathway and increased the glutathione peroxidase 4 (GPX4) level, thereby suppressing sorafenib-induced ferroptosis. The combination of sorafenib and RSL3, a GPX4 inhibitor, significantly inhibited GSTZ1 deficient cell viability and promoted ferroptosis, accompanied with ectopic increases of iron and lipid peroxides. An *in vivo* experiment showed that the combination of sorafenib and RSL3 had a synergic therapeutic effect on HCC progression in *Gstz1^−/−^* mice. In conclusion, GSTZ1 was significantly downregulated in sorafenib resistant hepatoma cells. GSTZ1 enhanced sorafenib-induced ferroptosis by inhibiting the NRF2/GPX4 axis in HCC cells. GSTZ1 deficiency was resistant to sorafenib-induced ferroptosis and is, therefore, a potential therapeutic approach for treating HCC by synergizing sorafenib and RSL3 to induce ferroptosis.

## Introduction

Hepatocellular carcinoma (HCC) is the fourth leading cause of cancer-related death worldwide ^1^. In the early stages of HCC, curative treatment can be achieved with tumor ablation, resection, or liver transplantation ^2^. However, majority of HCC patients are already in the middle-late stage when diagnosed; thus, the optimal period for curative treatment is missed. Sorafenib, a multi-target kinase inhibitor, has been confirmed to prolong the survival of advanced HCC patients to 6.5 months in phase III trial ^3^. Thus, it has been approved by the Food and Drug Agency as a first-line treatment for advanced HCC. However, several patients with advanced HCC have limited survival benefit due to acquired resistance to sorafenib, leading to a high recurrence rate ^4^. Therefore, the mechanism of sorafenib resistance needs to be explored, and new molecular targets should be identified.

Ferroptosis is a newly described programmed form of cell death characterized by iron-dependent accumulation of lipid peroxides to lethal amounts, different from the traditional cell death forms of apoptosis, necroptosis, and autophagy ^5^. Growing evidence indicates that ferroptosis can be induced by the inhibition of cystine/glutamate transporter (SLC7A11/xCT) activity, downregulation of glutathione peroxidase 4 (GPX4), and accumulation of iron and lipid reactive oxygen species (ROS) ^6–8^. Recent reports have shown that sorafenib could induce ferroptosis; thus, targeting ferroptosis to improve sorafenib therapy might be a new promising strategy for HCC treatment ^9–11^.

Glutathione S-transferases (GSTs) is a class of phase II detoxification enzymes that catalyze the conjugation of glutathione (GSH) to endogenous or exogenous electrophilic compounds ^12^. GSTs, including GSTM and GSTP ^13–15^, are involved in the development of chemotherapy resistance ^16,17^. Glutathione S-transferase zeta 1 (GSTZ1) is an important member of the GST superfamily. It participates in the catabolism of phenylalanine/tyrosine and catalyzes the isomerization of maleylacetoacetate to fumarylacetoacetate ^18^. We previously found that GSTZ1 was poorly expressed in HCC, and GSTZ1 deficiency could lead to metabolite succinylacetone accumulation and thereby activate the NRF2 signaling pathway ^19,20^. Considering the importance of GSTZ1 in the development and progression of HCC, GSTZ1 may be an anticancer hallmark for sorafenib resistance in HCC. Therefore, it is crucial to investigate the role of GSTZ1 in chemotherapeutic resistance and elucidate underlying mechanisms. In the present study, we investigated the role of GSTZ1 in sorafenib-induced ferroptosis in HCC cell lines *in vitro* and in Gstz1-knockout mice *in vivo*, and determined the involved molecular mechanisms. Our study not only identify a novel mechanism of sorafenib resistance but also suggest a new link between GSTZ1 and ferroptosis.

## Results

### GSTZ1 is downregulated in sorafenib-resistant HCC

To investigate the molecular mechanism of sorafenib resistance in HCC, we generated sorafenib-resistant (SR) HCC cell lines *in vitro*. Resistance was achieved by gradually increasing the concentration of sorafenib in the medium over repeated passages ^21^. Finally, resistant HepG2 and SNU449 cell lines were established. We confirmed the acquired resistance of these resistant cells named HepG2-SR and SUN449-SR toward sorafenib by comparing to the parental cells. The half maximal inhibitory concentrations (IC_50_) of HepG2-SR and SNU449-SR cells to sorafenib were 2-3 times higher than that of the parental cells at 17.09 μM and 15.43 μM respectively (Fig. 1A). In addition, we evaluated the cell viability of sensitive and resistant cells treated with sorafenib over a series of time points or at different concentrations for 24h and found that the SR cells became less sensitive to sorafenib (Fig. 1B-C). To verify the role of GSTZ1 in sorafenib-resistant HCC, we comprehensively analyzed the expression levels of GSTZ1 in HepG2 cells and HepG2 cells resistant to sorafenib in GSE62813 databases. The results showed that GSTZ1 was significantly downregulated in SR cells (Fig. 1D). Subsequently, we further validated the low levels of GSTZ1 expression in SR cell lines via quantitative reverse-transcription polymerase chain reaction (qRT-PCR) and Western blotting (Fig. 1E-F). Together, these data indicate that GSTZ1 may play a negative role in mediating the resistance to sorafenib in HCC cells.

**Fig. 1.**
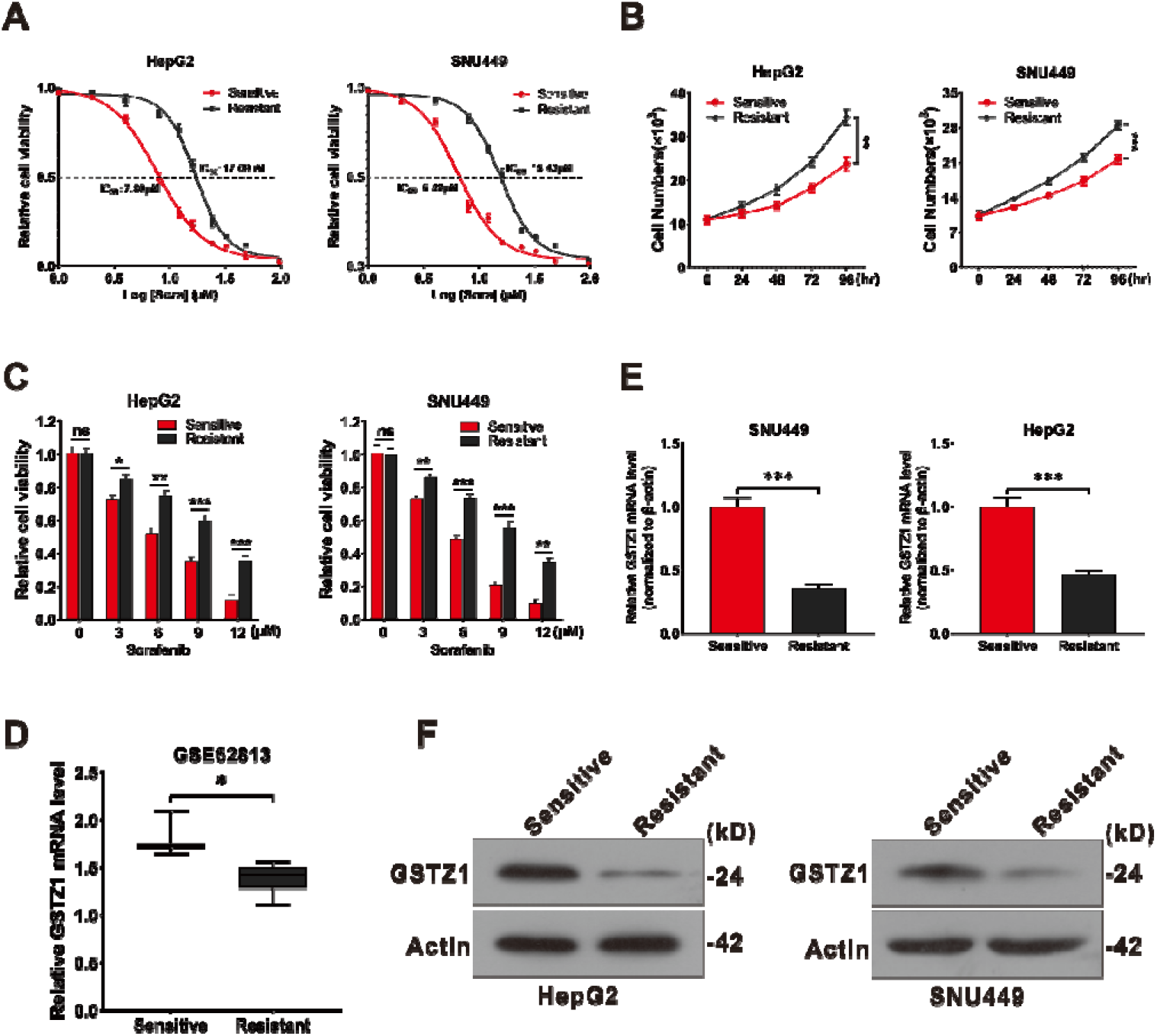
GSTZ1 is downregulated in sorafenib-resistant HCC cells. **a** The IC50 values of sorafenib-sensitive and sorafenib-resistant HCC cells treated with sorafenib. **b** Cell growth curve. **c** These sorafenib-sensitive and sorafenib-resistant HCCs (HepG2, SNU449) were treated with indicated concentrations of sorafenib for 24 h, and cell viability was assayed using the CCK-8 assay. **d** GSTZ1 RNA level in sorafenib-sensitive HepG2 cells (n=3) and sorafenib-resistant HepG2 cells (n=10). **e-f** mRNA and protein levels of GSTZ1 in sorafenib-sensitive and sorafenib-resistant cells. For Western blotting, 50 μg protein was loaded per well. HCC: hepatocellular carcinoma. Values represent the mean ± standard deviation (SD) (n = 3, performed in triplicate). ns: no significant difference, *p < 0.05, **p < 0.01, ^***^p < 0.001, Student’s *t*-test (two groups) or one-way ANOVA followed by Tukey tests (three groups).

### GSTZ1 Knockout promotes sorafenib resistance in HCC

To further evaluate whether GSTZ1 is related to sorafenib-resistance in HCC, we found that overexpression of GSTZ1 through an adenovirus system ^19,20^ increased the sensitivity of HCC cells to sorafenib and inhibited cell proliferation via morphological observation. Conversely, knockout of GSTZ1 in HepG2 and SNU449 cells via the CRISPR-Cas9 system ^19,20^ decreased the drug sensitivity and weakened the growth inhibition effect of sorafenib (Fig. 2A-B). Next, we analyzed the cell viability of GSTZ1 overexpression (OE) and knockout (KO) cells treated with sorafenib over a series of time points or at different concentrations for 24 h by cell growth curve. As expected, GSTZ1 overexpression significantly enhanced the sensitivity of HCC cell lines to sorafenib (Fig. 2C-D). No surprisingly, the IC_50_ value of GSTZ1-OE cells was decreased compared to that of the control groups, whereas the IC_50_ value of GSTZ1-KO groups was increased (Fig. 2E-F). Taken all together, our results showed that GSTZ1 deficiency enhanced sorafenib resistance in HCC.

**Fig. 2.**
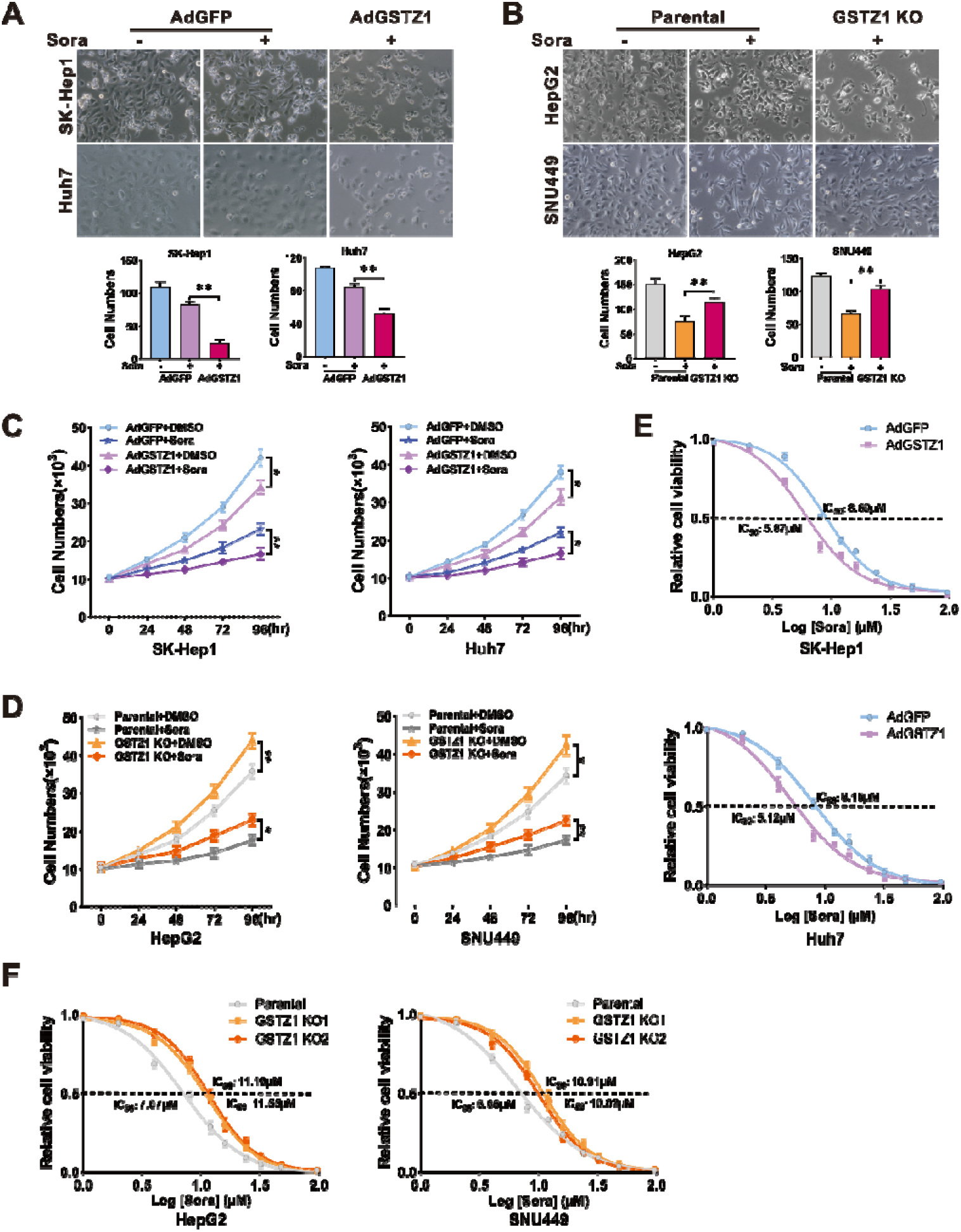
GSTZ1 knockout promotes sorafenib resistance in HCC cells. **a-b** Morphological phase-contrast images (top) and quantification (bottom) of cells after treatment with or without sorafenib (10 μM) for 24 h. Magnifications: ×200. **c-d** Cell growth curve. GSTZ1 overexpression (OE) (**c**) and GSTZ1 knockout (KO) (**d**) cells were treated with or without sorafenib (10 μM). **e-f** The IC50 of GSTZ1-OE (**g**) and GSTZ1-KO (**h**) cells were determined using the CCK-8 assay. HCC: hepatocellular carcinoma, DMSO: dimethyl sulphoxide, Sora: sorafenib. Values represent the mean ± SD (n = 3, performed in triplicate). *p < 0.05, **p < 0.01, Student’s t-test (two groups) or one-way ANOVA followed by Tukey tests (three groups).

### GSTZ1 overexpression enhances sorafenib-induced ferroptosis in HCC

Recent studies indicate that ferroptosis plays a key role in the chemoresistance of human cancers ^22–24^. We confirmed that sorafenib-induced cell death in HCC cell lines was blocked by ferrostatin-1 (Fer-1, an inhibitor of ferroptosis), deferoxamine (DFO, an iron chelator), and N-acetyl-L-cysteine (NAC, an antioxidant), but not by bafilomycin A1 (Baf-A1, an inhibitor of autophagy), ZVAD-FMK, and necrosulfonamide (Nec, an inhibitor of necroptosis). This suggested that ferroptosis, rather than apoptosis, is essential for sorafenib-induced cell death in HCC (Supplementary Fig. 1A), consistent with previous studies ^11,25,26^. To determine whether GSTZ1 played a role in ferroptosis to reduce sorafenib resistance in HCC, transmission electron microscopy (TEM) was used to observe the morphological changes in sorafenib-induced HCC cells with or without GSTZ1 depletion. GSTZ1-OE cells treated with sorafenib displayed smaller mitochondria, diminished or vanished mitochondria crista, and condensed mitochondrial membrane densities compared to parental cells, whereas GSTZ1-KO alleviated the abnormalities of mitochondrial morphology and cell death induced by sorafenib (Fig. 3A). To further verify this observation, we measured ROS, iron and lipid peroxidation levels, which are the primary cause of ferroptosis^27^, after interference with GSTZ1 expression. Results showed that GSTZ1 overexpression significantly increased ROS, iron, and MDA level accumulation in sorafenib-induced HCC cell lines (Fig. 3B-D and Supplementary Fig. 1B-C), whereas GSTZ1 knockout decreased their levels. In addition, we assessed mRNA and protein expression levels of ferroptosis-associated genes. Results showed that GSTZ1-OE decreased the expression levels of ferroptosis-related genes in sorafenib or erastin (an inducer of ferroptosis)-induced hepatoma cells, including *GPX4*, *FTL*, and *SLC7A11*. In contrast, GSTZ1-KO increased the levels of these above genes (Fig. 3E-G and Supplementary Fig. 1D-E). Interestingly, ferroptosis-associated genes were also enhanced in HepG2-SR and SNU449-SR cells (Fig. 4A-B), which had relatively lower levels of GSTZ1 than the parental cells. Meanwhile, GSTZ1 overexpression in drug-resistant cells reduced the levels of ferroptosis-related genes (Fig. 4C), increased the accumulation of iron and MDA levels, and enhanced the inhibition of sorafenib to resistant cells (Fig. 4D-F), consistent with the sorafenib-sensitive cells. To further identify the role of ferroptosis in sorafenib resistance caused by GSTZ1 deficiency, we examined the curative effects of sorafenib by intervention of ferroptosis. We observed that ferrostatin-1 inhibited sorafenib-induced GSTZ1-OE cell death, whereas erastin promoted GSTZ1-KO cell death (Fig. 4G). These results suggested that GSTZ1 increased the sensitivity of hepatoma cells to sorafenib by inducing ferroptosis.

**Fig. 3.**
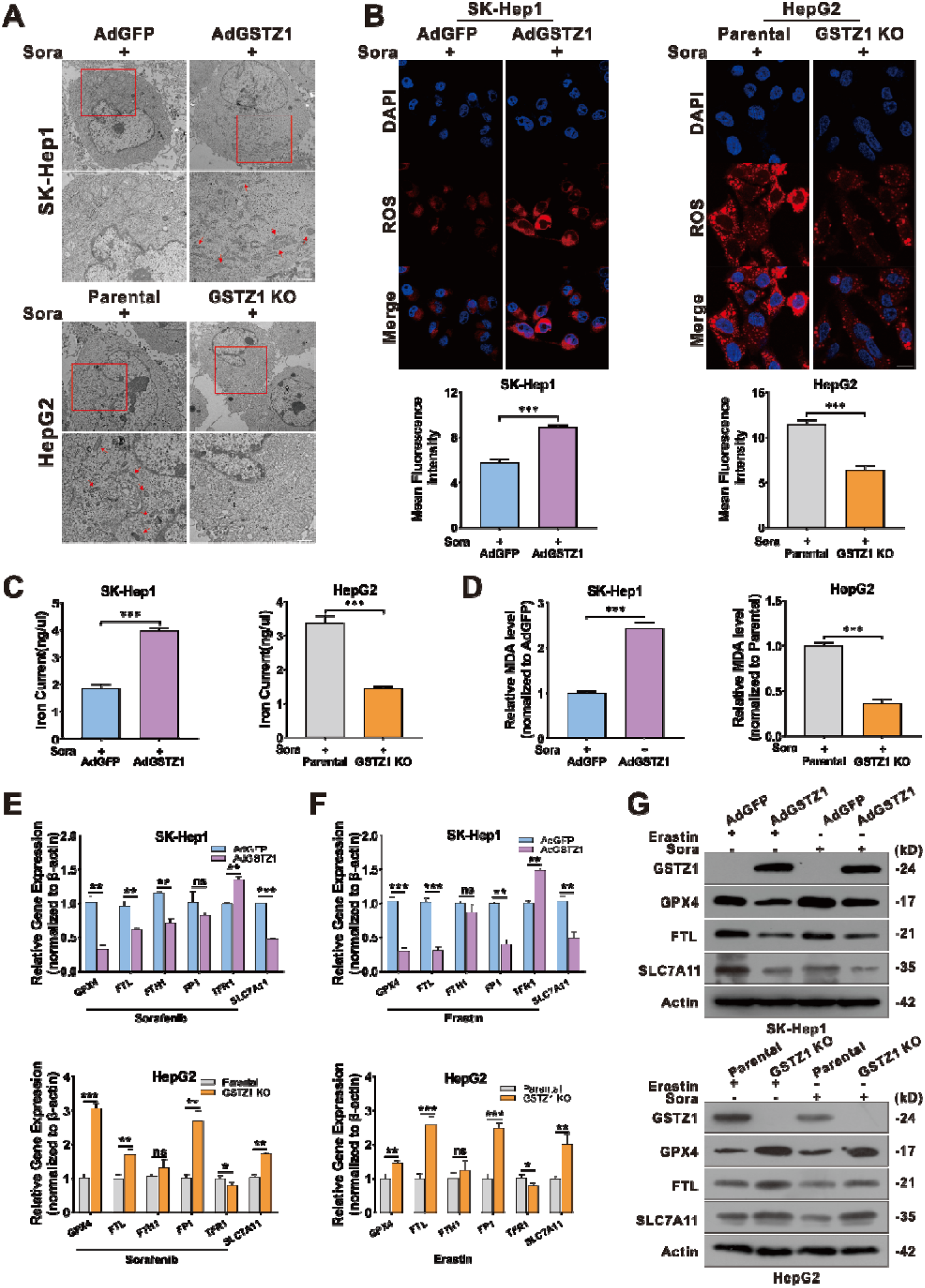
GSTZ1 overexpression enhances sorafenib-induced ferroptosis in HCC. **a** Representative TEM images of the mitochondrial morphology in GSTZ1-OE SK-Hep1 and GSTZ1-KO HepG2 cells treated with 10 μM sorafenib for 24 h. Red arrows indicate mitochondria. Bar = 1 μm. **b** Representative images (top) and quantification (bottom) of ROS level in GSTZ1-OE and GSTZ1-KO cells treated with sorafenib for 24 h. Bar = 20 μm. **c-d** The intracellular iron (**c**) and MDA (**d**) levels in GSTZ1-OE and GSTZ1-KO cells treated with sorafenib for 24 h. **e-g** mRNA and protein levels of target genes associated with ferroptosis in GSTZ1-OE and GSTZ1-KO cells treated with sorafenib or erastin, determined via qRT-PCR (**e-f**) and Western blotting (**g**), respectively. For Western blotting, 50 μg protein was loaded per well. HCC: hepatocellular carcinoma, Sora: sorafenib, ROS: reactive oxygen species, MDA: malondialdehyde. Values represent the mean ± SD (n = 3, performed in triplicate). ns: no significant difference, *p < 0.05, **p < 0.01, ***p < 0.001, Student’s *t*-test (two groups) or one-way ANOVA followed by Tukey tests (three groups).

**Fig. 4.**
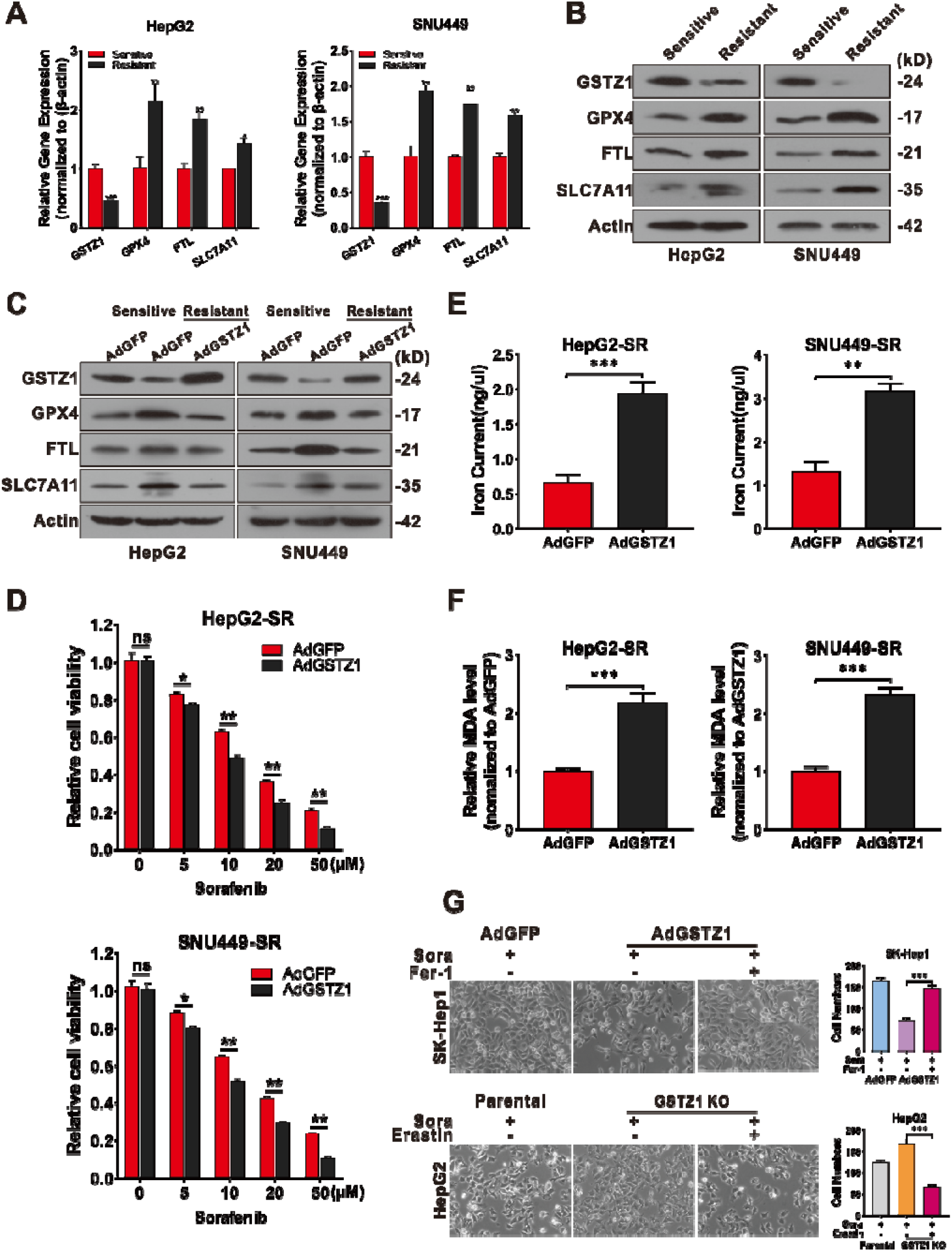
GSTZ1 overexpression sensitizes hepatoma cells to sorafenib-induced ferroptosis. **a-b** The mRNA (**a**) and protein (**b**) levels of ferroptosis-related genes in sorafenib-sensitive and sorafenib-resistant cells were assayed using qRT-PCR and Western blotting, respectively. **c** Western blotting for assessment of protein levels of ferroptosis-related genes in sorafenib-resistant HCCs with adenoviruses expressing GFP (AdGFP) or GSTZ1 (AdGSTZ1). **d** The cell viability of sorafenib-resistant cell with GSTZ1 overexpression was determined using CCK-8 assay. **e-f** The iron (**e**) and MDA (**f**) levels in GSTZ1-OE sorafenib-resistant cells. **g** The morphology (left) and quantification (right) of indicated HCC cells treated with sorafenib (10 μM for 24 h) alone or in combination with Fer-1 (1 μM for 24 h) or erastin (10 μM for 24 h). Magnifications: ×200. For Western blotting, 50 μg protein was loaded per well. HCC: hepatocellular carcinoma, Sora: sorafenib, MDA: malondialdehyde, Fer-1: ferrostatin-1, SR: sorafenib resistant. Values represent the mean ± SD (n = 3, performed in triplicate). ns: no significant difference, *p < 0.05, **p < 0.01, ***p < 0.001, Student’s t-test (two groups) or one-way ANOVA followed by Tukey tests (three groups).

### GSTZ1 sensitizes hepatoma cells to sorafenib-induced ferroptosis through the NRF2 signaling pathway

Previous studies have shown that GSTZ1 deficiency activated NRF2 pathway ^19,20^. Activation of NRF2 pathway plays a critical role in protecting HCC cells against sorafenib-induced ferroptosis ^26^. To verify whether GSTZ1 regulated sorafenib-induced ferroptosis through the NRF2 signaling pathway, we blocked the NRF2 pathway using brusatol (an inhibitor of NRF2) or Flag-tagged Kelch-like ECH-associated protein 1 (KEAP1) ^28^ (an cytosolic inhibitor of NRF2) in GSTZ1-KO cells and observed the characteristic indicators related to ferroptosis, including MDA, iron, ROS, and 4-HNE levels. Unexpectedly, the results demonstrated that NRF2 inhibition significantly increased the accumulation of these indicators in GSTZ1-KO cells (Fig. 5A-D right, Fig. 5E-F bottom, and Supplementary Fig. 2A-C bottom), whereas NRF2 activation using tertiary butylhydroquinone (tBHQ, an activator of NRF2) and Myc-tagged NRF2 yielded opposite results in GSTZ1-OE cells (Fig. 5A-D left, Fig. 5E-F top and Supplementary Fig. 2A-C top). Moreover, the morphological images also indicated that NRF2 inhibition increased the efficiency of sorafenib for growth inhibition in GSTZ1-depleted HCCs, whereas NRF2 activation decreased that in GSTZ1-OE cells (Fig. 6A). Importantly, the protein levels of ferroptosis-related genes were changed accordingly in GSTZ1-OE and −KO cells when treated with tBHQ or brusatol via Western blotting (Fig. 6B and Supplementary Fig. 2D). The above data suggested that GSTZ1 depletion alleviated sorafenib-induced ferroptosis via activation of the NRF2 pathway. As GPX4 is involved in ferroptosis and a transcriptional target gene of NRF2 ^29,30^, we utilized RSL3 to further examine whether ferroptosis is involved in the sensitivity of HCC cells to sorafenib. Interestingly, GPX4 inactivation enhanced sorafenib-induced ferroptosis and inhibited cell growth in GSTZ1-KO (Fig. 6C-E) and SR cells (Fig. 6F-H). Collectively, these findings indicated that the inhibition of NRF2 could markedly sensitize GSTZ1-deficient hepatoma cells to sorafenib treatment. Furthermore, we further examined targeting GPX4 could also improve the response of hepatoma cells to sorafenib.

**Fig. 5.**
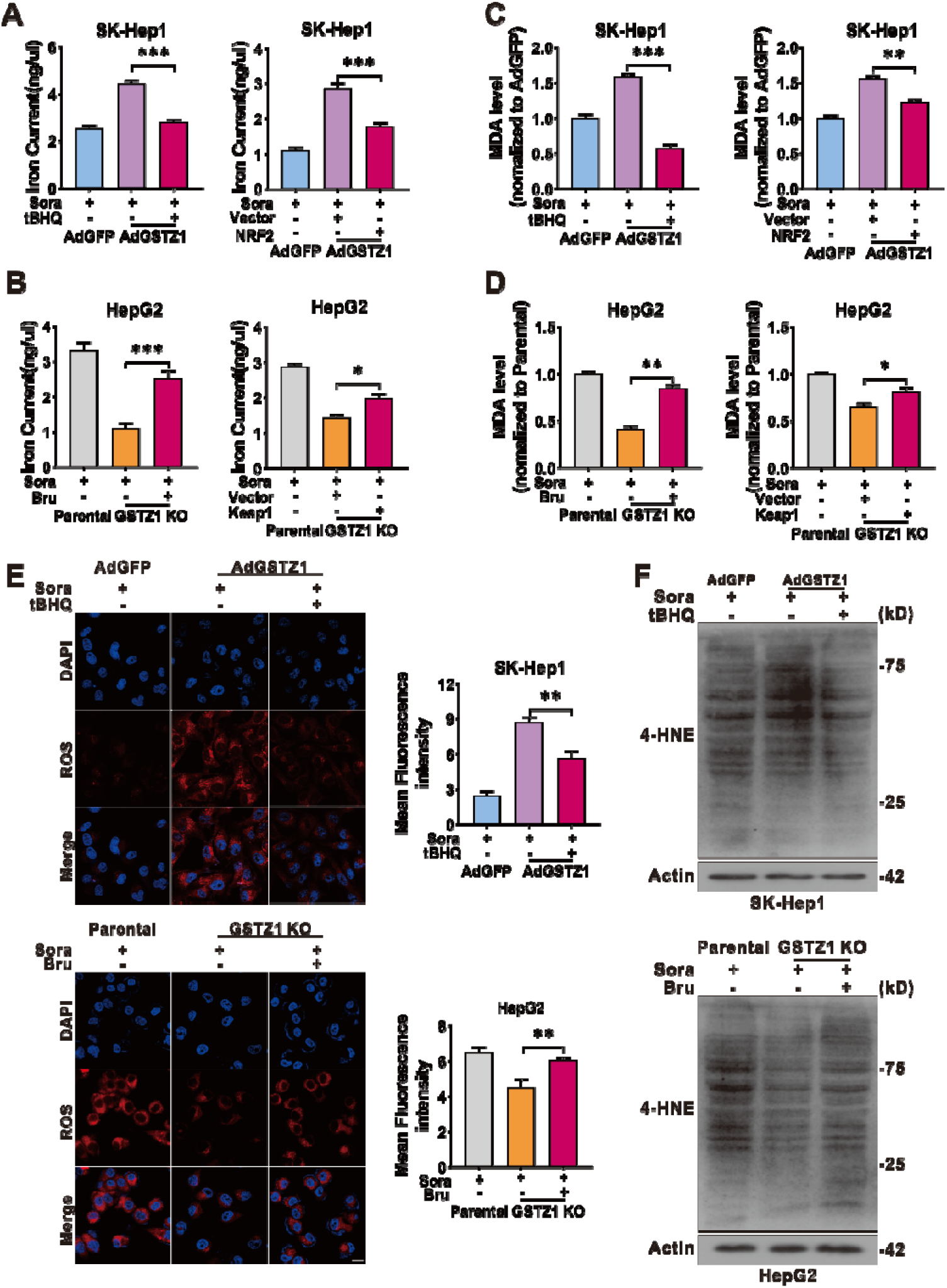
GSTZ1 knockout cells are insensitive to sorafenib-induced ferroptosis through the activation of NRF2. **a** GSTZ1-OE cells were treated with sorafenib alone or in combination with tBHQ (100 μM for 3 h) (left). GSTZ1-KO cells were treated with sorafenib alone or in combination with Bru (40 nM for 24 h) (left). Expressing Flag-KEAP1 plasmid was transfected into GSTZ1-OE cells with sorafenib treatment (right). Expressing Myc-NRF2 plasmid was transfected into GSTZ1-KO cells with sorafenib treatment (right). Levels of iron (**a-b**), and MDA (**c-d**) in these cells were assayed. **e** Representative images (top) and quantification (bottom) of ROS level in GSTZ1-OE cells treated with sorafenib alone or in combination with tBHQ (top) and GSTZ1-KO cells treated with sorafenib alone or in combination with Bru (bottom). Bar = 20 μm. **f** 4-HNE-induced protein modification were examined. The cell processing is described as above. For Western blotting, 50 μg protein was loaded per well. tBHQ: tertiary butylhydroquinone, Bru: brusatol, Sora: sorafenib, MDA: malondialdehyde, 4-HNE: 4-hydroxy-2-nonenal. Values represent the mean ± SD (n = 3, performed in triplicate). ns: no significant difference, *p < 0.05, **p < 0.01, ***p < 0.001, Student’s t-test (two groups) or one-way ANOVA followed by Tukey tests (three groups).

**Fig. 6.**
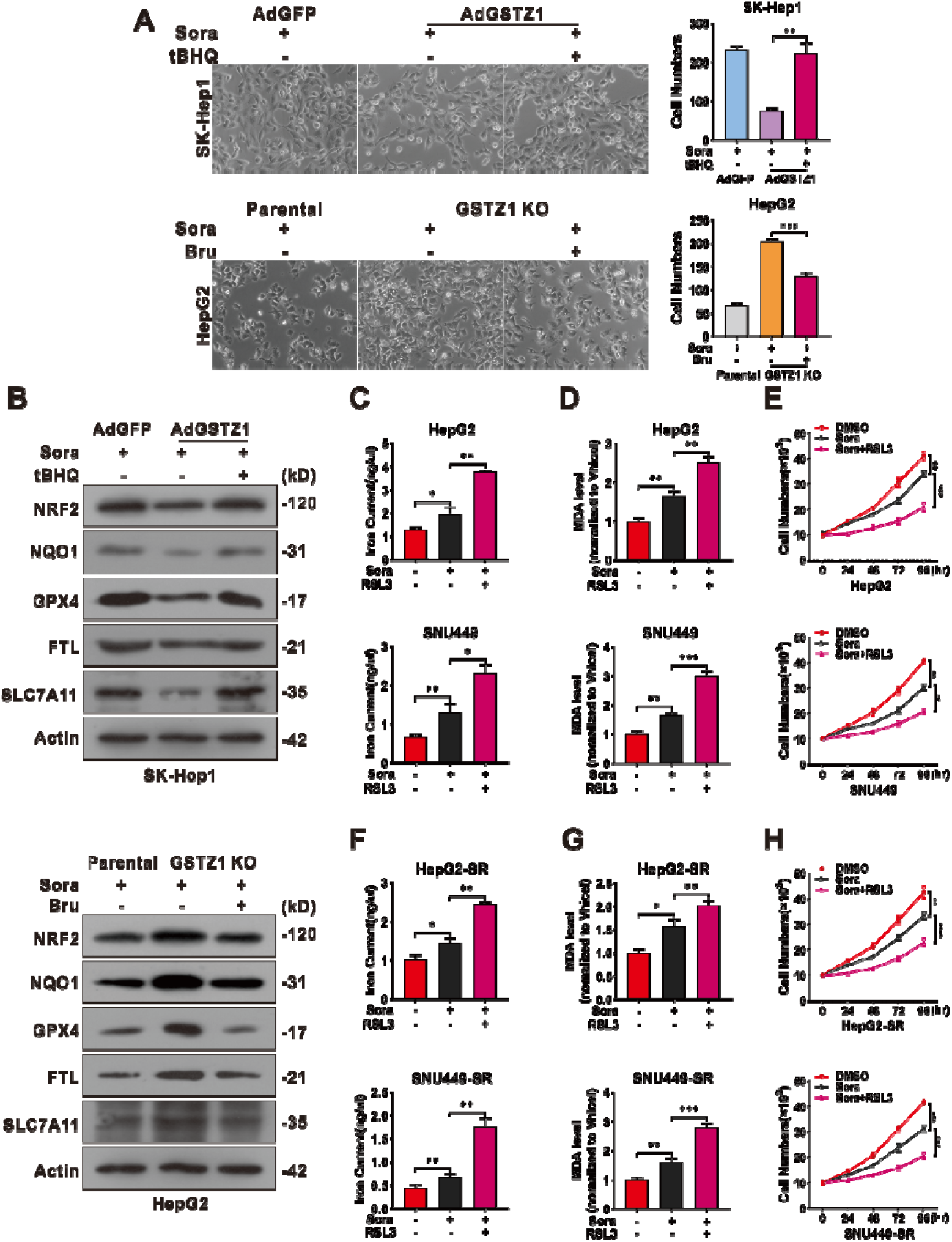
RSL3 enhances the sensitivity of GSTZ1-KO and sorafenib-resistant cells to sorafenib. **a-b** Morphological changes (**a**) and protein level (**b**) of ferroptosis-related genes in GSTZ1-OE SK-Hep1 cells treated with sorafenib alone or in combination with tBHQ (top) and GSTZ1-KO HepG2 cells treated with sorafenib alone or in combination with Bru (bottom). Magnifications: ×200. **c-d** The iron (left) and MDA (right) levels in GSTZ1-KO cells treated with sorafenib alone or in combination with RSL3 (500nM for 24 h). **e** The cell growth curve of GSTZ1-KO cells treated with sorafenib alone or in combination with RSL3. **f-g** The iron (left) and MDA (right) levels in sorafenib-resistant cells treated with sorafenib alone or in combination with RSL3. **h** The cell growth curve of sorafenib-resistant cells treated with sorafenib alone or in combination with RSL3. For Western blotting, 50 μg protein was loaded per well. RSL3: Ras-selective lethal small molecule 3, tBHQ: tertiary butylhydroquinone, Bru: brusatol, DMSO: dimethyl sulphoxide, Sora: sorafenib, MDA: malondialdehyde. Values represent the mean ± SD (n = 3, performed in triplicate). *p < 0.05, **p < 0.01, ***p < 0.001, Student’s t-test (two groups) or one-way ANOVA followed by Tukey tests (three groups).

### RSL3 enhances the anticancer activity of sorafenib in *Gstz1*^−/−^ mice

To further investigated the role of GSTZ1 in mediating sorafenib resistance in HCC progression *in vivo*, we established the mouse model of liver cancer induced by DEN/CCl_4_ as our previous induction method ^20^ and drug administration with three regimens: DMSO, sorafenib (30mg/kg, every 2 days for 4 weeks), RSL3 (10mg/kg, every 2 days for 4 weeks) (Fig. 7A). Compared with wild type (WT) mice, *Gstz1* knockout significantly reduced the inhibitory effects of sorafenib treatment *in vivo* than that in WT mice, as indicated by the increased tumor sizes and number of tumor nodules and higher level of alanine aminotransferase (ALT) in serum (Fig. 7B-E). Moreover, sorafenib combined with RSL3 had a more significant protective effect on tumorigenesis in *Gstz1^−/−^* mice than sorafenib alone. To substantiate the role of GSTZ1 in regulating ferroptosis-mediated sorafenib resistance *in vivo*, we detected the levels of iron, 4-HNE modification, MDA, ROS and ferroptosis-associated gene expression mRNA and protein in the liver tumor tissues. Consistent with the results *in vitro*, *Gstz1* knockout decreased the sensitivity of HCC to sorafenib by weakening ferroptosis. Meanwhile, RSL3/sorafenib combination treatment reduced drug resistance caused by GSTZ1 depletion (Fig. 7F-J and Supplementary Fig. 3A). Furthermore, histological analysis indicated that the cytoplasm and nuclei of liver tumors in the combination treatment groups exhibited weaker immunoreactivity for GPX4 and Ki67 respectively than that in sorafenib-alone groups. RSL3 significantly enhanced the inhibitory effects of sorafenib on cell proliferation in *Gstz1^−/−^* mice, which was highly consistent with the in vitro results (Fig. 7K). These results indicate that targeting the NRF2/GPX4 axis using RSL3 significantly enhances sorafenib-induced ferroptosis and inhibits hepatocarcinogenesis *in vivo*.

**Fig. 7.**
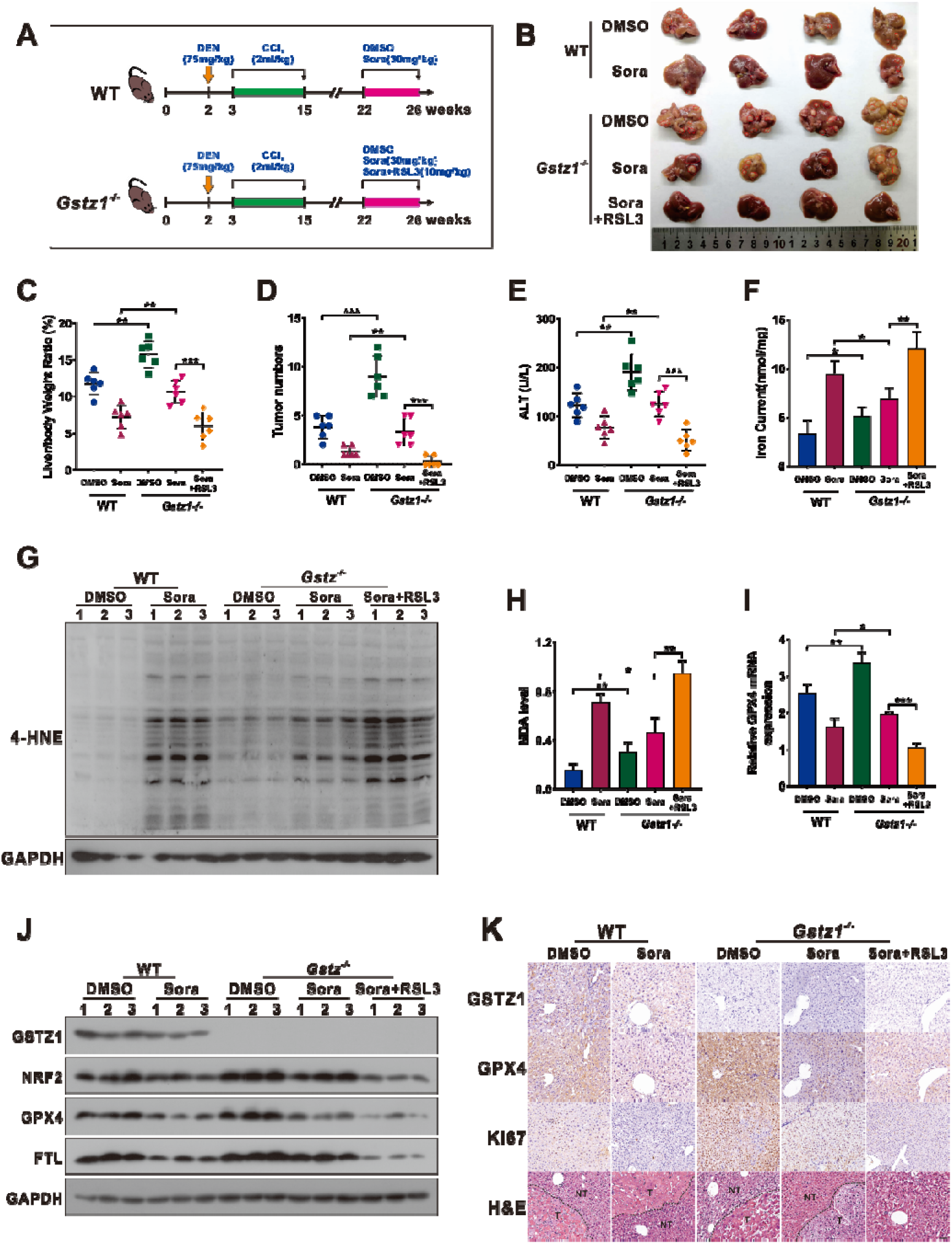
RSL3 enhances the anticancer activity of sorafenib in *Gstz1^−/−^* mice. **a** Schematic representation of the experimental design for mice. **b** Gross appearances of liver tumors. The red circles represent tumors. **c-e** In vivo analyses of liver/body weight ratio (**c**), tumor numbers (**d**), and serum alanine aminotransferase (ALT) (**e**) levels of the five groups. **f-h** The levels of iron (**f**) and MDA (**h**) in mice were assayed. Western blotting to assess 4-HNE modification level (**g**) in murine livers. **i-j** mRNA (**i**) and protein (**j**) levels of GPX4, FTL, and SLC7A11 in the five groups of liver tumors as assessed using Western blotting and real-time qPCR, respectively. **k** Representative H&E staining and immunohistochemistry images of GSTZ1, GPX4, and Ki67 in hepatic tumors. Bar = 50 μm. For Western blotting, 50 μg protein was loaded per well. WT: wild type, DEN: diethylnitrosamine, CCl_4_: carbon tetrachloride, DMSO: dimethyl sulphoxide, Sora: sorafenib, RSL3: Ras-selective lethal small molecule 3, ALT: alanine aminotransferase, 4-HNE: 4-hydroxy-2-nonenal, H&E: hematoxylin and eosin. Values represent the mean ± SD (n = 3, performed in triplicate). *p < 0.05, **p < 0.01, ***p < 0.001, Student’s t-test (two groups) or one-way ANOVA followed by Tukey tests (five groups).

## Discussion

The incidence of HCC continues to increase globally, and HCC remains to have high incidence and mortality rates ^31^. Sorafenib resistance remains a treatment challenge in HCC and leads to poor prognosis ^32^. Therefore, the comprehensive elucidation of the underlying mechanism of sorafenib resistance in HCC may improve the curative effect of chemotherapy and guide the clinical medication. Herein, we found that GSTZ1 depletion activated the NRF2/GPX4 pathway and inhibited sorafenib-induced cell death, accompanied by the compromised accumulation of iron level, lipid peroxidation, and subsequent ferroptosis. Hence, blocking the NRF2/GPX4 pathway to enhance the anticancer activity of sorafenib by inducing ferroptosis represents a promising therapeutic strategy for the treatment of HCC (Fig. 8).

**Fig. 8.**
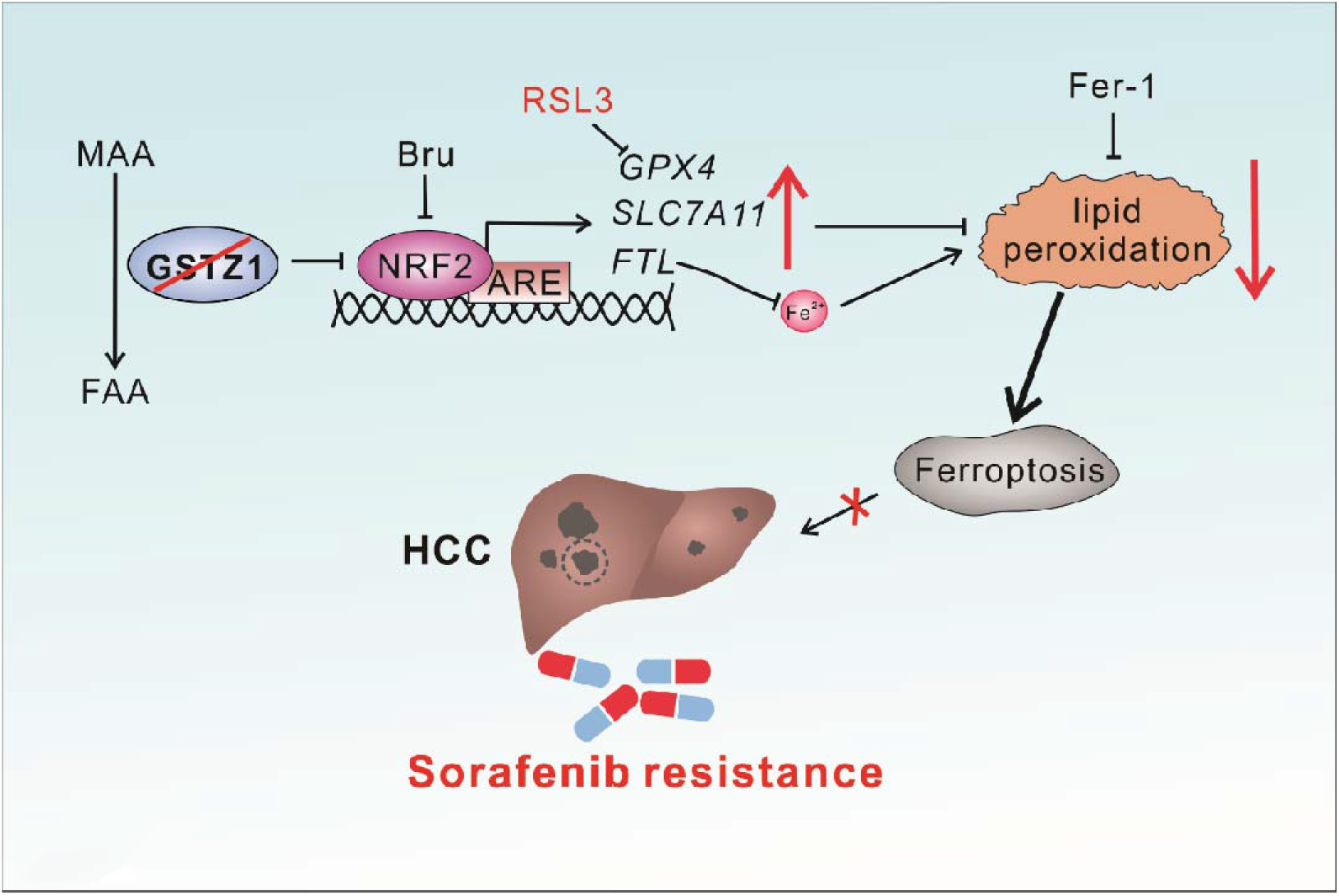
A proposed model of the resistance of GSTZ1-deficient cells to sorafenib. MAA: maleylacetoacetate, FAA: fumarylacetoacetate, Bru: brusatol, ARE: anti-oxidation response element. Fer-1: ferrostatin-1, HCC: hepatocellular carcinoma.

GSTs are phase II detoxification enzymes that play important roles in protecting cellular macromolecules from both oxidative stress and carcinogenic electrophiles ^33^. The major roles of GSTs in the detoxification of xenobiotics predicts their important role in drug resistance. Tumor cells may develop resistance to alkylating anticancer drugs by increasing the levels of GSTs ^34^. Several subclasses in the GST family contribute to chemoresistance in various cancers ^15,35–40^. GSTM1 was reported to be a predictive biomarker for the efficacy of chemotherapy in breast cancer and ovarian cancer ^15,35^.

GSTP expression was also correlated with platinum drug chemosensitivity and prognosis in ovarian cancer, pancreatic ductal adenocarcinoma, and lung cancer ^36–39^. As a member of the GST family, GSTZ1 plays a similar detoxification role, but it is independently characterized as a maleylacetoacetate isomerase (MAAI), which is essential for phenylalanine metabolism ^18^. We previously reported that GSTZ1 is downregulated in HCC, leading to increased accumulation of the carcinogenic metabolite succinylacetone and activation of the NRF2/IGFIR pathways through inactivation of KEAP1 ^19^. The current study demonstrates that GSTZ1 is also downregulated in sorafenib-resistant HCC cells. Furthermore, re-expression of GSTZ1 enhances the sensitivity of HCC cells to sorafenib treatment, indicating the negative role of GSTZ1 in sorafenib resistance.

Iron is an essential element for the synthesis of iron sulfur clusters, serving an important role in numerous cellular processes ^41^. Cancer cells exhibit a higher dependence on iron than normal cells ^42^, making them more susceptible to iron-catalyzed necrosis. This form of cell death was first defined as ferroptosis in 2012 ^5^, which characterized by the accumulation of lipid peroxidation products and lethal ROS derived from iron metabolism ^6,27^. An increasing number of small molecule compounds (e.g., erastin) or clinical drugs (e.g., sulfasalazine) has been found to induce ferroptosis by modulating iron metabolism and enhancing the accumulation of lipid peroxidation ^6,43^. As a homeostatic dysfunction of ferroptosis is believed to be an essential cause of chemoresistance ^44^, it is crucial to explore how to enhance the sensitivity of cancer cells to clinical chemotherapy drugs by triggering ferroptosis. In the case, ferroptosis inducer erastin significantly enhances the anticancer activity of cytarabine and doxorubicin in leukemia cells ^45^. It also reverses the resistance of ovarian cancer cells to cisplatin ^46^. Multiple studies recently verified that sorafenib plays an important role in inducing ferroptosis. Herein, we found that GSTZ1 deficiency aggravated the resistance to sorafenib-induced ferroptosis by preventing iron accumulation and lipid peroxidation production and decreasing the ROS level. In contrast, GSTZ1 overexpression increased the sensitivity of HCC cells to sorafenib by facilitating ferroptosis *in vitro*. Previous studies found that sorafenib-induced hepatoma cell death is mainly dependent on triggering ferroptosis by inhibiting of SLC7A11/xCT ^10,11,24^, and consistent findings were observed in this study.

As such, inducing ferroptosis may be a promising strategy for enhancing the sensitivity of tumor cells to chemotherapy. Haloperidol, a sigma-1 receptor (S1R) antagonist, promotes sorafenib-induced ferroptotic death by increasing ROS accumulation ^47^. Meanwhile, metallothionein-1G silencing (MT-1G) was reported to enhance the sensitivity of hepatoma cells to sorafenib by triggering ferroptosis ^24^. Collectively, our findings and those of previous studies suggest that ferroptosis plays an important role in the anti-tumor efficacy of sorafenib. Further, our data indicated that GSTZ1 plays a positive regulatory role in ferroptosis during sorafenib treatment.

Changes in certain metabolic pathways are also involved in the regulation of cell sensitivity to ferroptosis, including coenzyme Q10 consumption ^48^, decreased intracellular reducer such as NAPDH ^49^, and altered iron metabolism ^50^. Many components of the ferroptosis cascade are target genes of the transcription factor NRF2, indicating the critical role of the NRF2 pathway in mediating ferroptotic response ^29,30,51^. For example, the inhibition of p62-Keap1-NRF2 pathway significantly enhanced the anticancer activity of erastin and sorafenib by inducing ferroptosis in HCC cells *in vitro* and *in vivo* [27]. Consistent with previous reports, our results demonstrated that GSTZ1 deficiency markedly reduces sorafenib-induced ferroptotic cell death by increasing the level of NRF2 and ferroptosis-related genes including GPX4, SLC7A11, and FTL. In contrast, pharmacological- or Keap1-mediated inhibition of NRF2 increases the sensitivity of GSTZ1-deficient cells to sorafenib by enhancing ferroptosis *in vitro*. Meanwhile, GPX4 is the only reported enzyme that is capable of directly reducing complex phospholipid hydroperoxides and is a downstream target gene of NRF2. Therefore, targeting GPX4 is currently considered to be a crucial strategy for triggering ferroptosis ^7,44^ Mechanistically, we verified that GSTZ1 knockout inhibited sorafenib-induced ferroptosis by activation of the NRF2/GPX4 axis *in vitro* and *in vivo*. Moreover, targeting GPX4 using RSL3 in GSTZ1-knockout and sorafenib-resistant HCC cells significantly increased iron accumulation, ROS level, and lipid peroxidation production and enhanced sorafenib-induced inhibition of cell proliferation. Importantly, GPX4 inhibition using RSL3 with sorafenib therapy elicited a significant tumor regression in *Gstz1^−/−^* mouse models *in vivo*.

To our best knowledge, this is the first study to explore the role of GSTZ1 in sorafenib resistance in HCC. Our findings provide new insights into the molecular basis of the role of GSTZ1 in sorafenib resistance, and indicate that sorafenib combined with RSL3 can synergistically overcome acquired resistance to sorafenib and improve the anticancer efficacy of sorafenib in HCC. Blocking the NRF2/GPX4 axis may have a therapeutic benefit in HCC patients with GSTZ1 deficiency. Our findings also demonstrate the sensitizing role of RSL3 for enhancing sorafenib effectiveness. Importantly, GSTZ1 deficiency was resistant to sorafenib-induced ferroptosis and is therefore a potential therapeutic approach for treating HCC by synergizing sorafenib and RSL3 to induce ferroptosis.

## Materials and Methods

### Cell lines

Human hepatoma cell lines SK-Hep1, HepG2 and SNU449 were directly obtained from American Type Culture Collection (ATCC, VA, USA). Huh7 cells were obtained from Cell Bank of the Chinese Academy of Sciences (Shanghai, China). These cells were cultured in Dulbecco’s modified Eagle’s medium (SK-Hep1, HepG2, Huh7) or RPMI 1640 medium (SNU449) supplemented with 10% fetal bovine serum (FBS; Gibco, Rockville, MD, USA), 100units/mL penicillin and 100mg/mL streptomycin in a humidified incubator at 37 °C containing 5% CO_2_.

### Reagents and antibodies

Erastin (HY-15763), Ferrostatin-1 (HY-100579), Deferoxamine (HY-B0988) and Necrosulfonamide (HY-100573) were purchased from MedChemExpress (MCE; Shanghai, China). Sorafenib (S7397), ZVAD-FMK (S7023), Bafilomycin A1 (S1413) and RSL3 (S8155) were obtained from Selleckchem (Houston, TX, USA). N-acetyl-L-cysteine (NAC, S0077) was from Beyotime (Shanghai, China). Brusatol (Bru, MB7292) was obtained from Meilunbio (Dalian, China). Tertiary butylhydroquinone (tBHQ, 112941) was obtained from Sigma (Shanghai, China). Antibodies raised against GPX4 (ab125066), NRF2 (ab62352), 4-HNE(ab46545), NQO1(ab34173) and β-actin (ab6276) were obtained from Abcam (Cambridge, MA, USA), anti-SLC7A11 (NB300-318) was from Novusbio (Centennial, CO, USA), anti-FTL (10727-1-AP) was from Proteintech (Shanghai, China), and anti-GSTZ1 (GTX106109) was from GeneTex (San Antonio, CA, USA).

### Generation of sorafenib-resistant cell lines

To establish sorafenib-resistant cells, HepG2 and SNU449 cells were cultured by exposing cells with sorafenib at 5% of IC50 concentration and the concentration was gradually increased at 10% of IC50 until the maximum tolerated doses (10 μM) have been reached. Sorafenib-resistant cells (HepG2-SR and SNU449-SR) were cultured continuously at 1 μM concentration of sorafenib to maintain the acquired resistance.

### Quantitative real-time polymerase chain reaction (qRT-PCR)

Total RNA was isolated from HCC cell lines using TRIzol reagent (Invitrogen, Rockville, MD, USA) according to the manufacturer’s instructions. Purified RNA samples were reverse-transcribed into cDNA using the PrimeScript™ RT Reagent Kit with gDNA Eraser (RR047A, TaKaRa, Tokyo, Japan).

Complementary DNA from cell samples was amplified with the specific primers (Supplementary Table. 1). Briefly, Real-time qPCR was performed to quantity mRNA levels, using the SYBR Green qPCR Master Mix (Bio-Rad, Hercules, CA, USA) in accordance with the manufacturer’s instructions. The objective CT values were normalized to that of β-actin and relative mRNA expression levels of genes were calculated using 2^−ΔΔCt^ method. Each sample was analyzed in triplicate.

### Western blot analysis

Protein samples from cells and animal tissues were lysed in Cell Lysis Buffer (Beyotime Biotechnology, Jiangsu, China) containing 1mM of phenylmethanesulfonyl fluoride (PMSF, Beyotime). The concentration of the protein homogenates was measured using the BCA protein assay Kit (Dingguo, Beijing, China). Equal volumes of protein samples were separated by SDS-poly acrylamide gel electrophoresis and electro-transferred to PVDF membranes (Millipore, Billerica, MA, USA). After blocked with 5% non-fat milk dissolved in TBST (10mM Tris, 150 mM NaCl, and 0.1% Tween-20; pH 7.6), for 2 h at room temperature, the membranes were incubated with the primary antibodies overnight at 4 °C. Thereafter, membranes were incubated with the secondary antibodies coupled to horseradish peroxidase (HRP) for 2 h at room temperature. Protein bands were visualized with enhanced Chemiluminescence substrate Kits (ECL, New Cell & Molecular Biotech Co, Ltd, China).

### Transmission electron microscope assay

Cells were collected and fixed with 2.5% glutaraldehyde. Subsequently, cells were postfixed in 2% osmium tetroxide and dehydrated through a series of graded ethyl alcohols. Samples were embedded in epoxy resin, sectioned, and placed onto nickel mesh grids. The images were acquired using a Hitachi-7500 transmission electron microscope (Hitachi, Tokyo, Japan).

### Intracellular ROS measurements

Cells were seeded on coverslips in a 12-well plate, and then treated with the varying concentrations of test compound or drug. After 24 h, cells were incubated at a final concentration of 5 μM CellROX® Orange reagent (Life Technologies, Carlsbad, USA) for 30 min at 37 °C, after which they were washed, dyed with DAPI, mounted with Anti-fade Mounting Medium, and immediately analyzed for fluorescence intensity under Leica Confocal Microscope (TCS SP8, Germany) with a 40× objective lens.

### Measurement of total iron contents in hepatoma cells and liver tissues

The iron concentration was assessed using the Iron Assay Kit (MAK025; Sigma) according to the manufacturer’s instructions. Briefly, tissues (10 mg) or cells (2 × 10^6^) were rapidly homogenized in 4-10 volumes of Iron Assay buffer. Tissue or cell homogenates was centrifuged at 16,000 × *g* for 10 minutes at 4 °C and removed insoluble material. To measure total iron, add 1-50 μL samples to sample wells in a 96 well plate, bring the volume to 100 μL per well with Iron Assay Buffer and add 5 μL Iron Reducer to each of the sample wells to reduce Fe^3+^ to Fe^2+^. And then samples were mixed using a horizontal shaker and incubated at 25 °C for 30 minutes. Subsequently, 100 μL Iron Probe were added and incubated the reaction for 1 hr at 25 °C. During each incubation, the plate was protected from light. Thereafter, the absorbance was detected at 593 nm using a microplate reader.

### Detection of malondialdehyde (MDA)

Analysis of lipid peroxidation was assessed by quantification of MDA concentration in cell lysates using a Lipid Peroxidation MDA Assay Kit (S0131) obtained from Beyotime in accordance with the manufacturer’s instructions.

### Cell growth curve and cell viability assay

For cell growth curve analysis, cells were seeded at 1×10^4^ cells/well in 96-well microtiter plates with three replicate per group and cultured overnight at 37°C in a humidified incubator containing 5% CO2. The plate was scanned and phase-contrast images were acquired after over a series of time points post treatment, and then quantified time-lapse curves were plotted using IncuCyte ZOOM software (Essen BioScience, Ann Arbor, MI, USA).

For cell viability assay, cells were seeded at 1,000 cells per well in 96-well plates with fresh medium and analysed by using the Cell Counting Kit-8 (CCK-8) (CK04, Dojindo, Japan) according to the manufacturer’s instructions. The microplates were incubated at 37°C for additional 2 h. Absorbance was read at 450 nm using a microplate reader (Thermo Fisher, USA).

### Half maximal inhibitory concentration assay (IC50)

The cells were planted in 96-well plates with fresh medium at 1.0×10^4^ cells per well. The corresponding concentrations of drug were given to cells for 24 h after the cultured plates were placed in a humidified incubator for 12 h. After 24 h, CCK-8 (Dojindo, Japan) was used to measure drug sensitivity at 450 nm using a microplate reader (Thermo Fisher, USA) after incubating at 37 °C for 1–2 h.

### Animal experiments

Heterozygous 129-*Gstz1*^tm1Jmfc^/Cnbc mice (EM: 04481) were purchased from the European Mouse Mutant Archive and were crossed to breed wild-type (WT) and *Gstz1^−/−^* mice. All mice were maintained under individual ventilation cages conditions in the laboratory animal center of Chongqing Medical University. For subsequent studies, mice were divided into five groups as follows: WT+DMSO (control), WT+Sora, *Gstz1^−/−^*+DMSO, *Gstz1^−/−^*+Sora, and *Gstz1^−/−^*+Sora+RSL3. Each group included three male and three female mice. At 2 weeks of age, all mice were administered an intraperitoneal injection of diethylnitrosamine (DEN; Sigma, St. Louis, MO, USA) at a dose of 75 mg/kg.

At the third week, the mice were intraperitoneally administered carbon tetrachloride (CCl_4_; Macklin, Shanghai, China) at 2 ml/kg twice a week for 12 weeks. In the WT+Sora and *Gstz1^−/−^*+Sora group, the mice at 22 weeks were administered intraperitoneally sorafenib (30mg/kg) every 2 days for 4 weeks until euthanasia. In the *Gstz1^−/−^*+Sora+RSL3 group, in addition to sorafenib administration as described above, the mice were injected intraperitoneally with RSL3 (10mg/kg) every 2 days for 4 weeks at the same weeks. Body weight of each mice was measured every week and retroorbital blood was collected before sacrifice. All mice were euthanized at 26 weeks of age. The liver weight and number of liver tumors were measured. Protein and mRNAs levels of hepatic tumors were detected by Western blotting and qRT-PCR analysis, respectively. The intrahepatic Iron and MDA levels were measured with Iron Assay Kit and MDA Assay Kit, respectively. Samples of liver tumor were collected for further study or fixed with 4% paraformaldehyde, embedded in paraffin, and sectioned for hematoxylin-eosin staining (H&E) and immunohistochemistry. All animal procedures were approved by the Research Ethics Committee of Chongqing Medical University (reference number: 2017010).

### Statistical analysis

All experiments were repeated independently with similar results at least three times. Statistical analysis and data plotting were performed using GraphPad Prism 7 (GraphPad Software, USA). All data were presented as mean ± standard deviation (SD) values. Unless mentioned otherwise, comparisons between two groups were performed by Student’s t-test, and Multiple-group comparisons were performed by the one-way ANOVA analysis with Scheffe post-hoc test. p < 0.05 was considered statistically significant.

## Acknowledgements

We wish to thank Dr. T.-C He (University of Chicago, USA) for providing the plasmids pAdEasy system, and Prof. Ding Xue (Tsinghua University) for supplying the CRISPR/Cas9 system.

## Authors’ contributions

ALH, NT and KW conceived the study and designed the experiments. QJW, BC and QX performed most experiments and analyzed the data. QZG assisted with experiments in *Gstz1^−/−^* knockout mice. QJW, NT and KW drafted and edited the manuscript with all authors providing feedback.

## Ethics Statement

This study was approved by the Research Ethics Committee of Chongqing Medical University (reference number: 2017010).

## Data Availability

The datasets used and/or analyzed during the current study are available from the corresponding author on reasonable request.

