## Supplementary for "GSTZ1 sensitizes hepatocellular carcinoma cells to sorafenib-induced ferroptosis via inhibition of NRF2/GPX4 axis"

Wang et al

**Contents**

**Supplementary Figure 1.** GSTZ1 overexpression enhances sorafenib-induced ferroptosis in HCC.

**Supplementary Figure 2.** GSTZ1 sensitizes hepatoma cells to sorafenib-induced ferroptosis through the NRF2 signaling pathway.

**Supplementary Figure 3.** RSL3 enhances the anticancer activity of sorafenib in *Gstz1*-knockout mice.

**Supplementary Table 1.** Primer sequences used in this study.

### Supplementary figures, table and legends

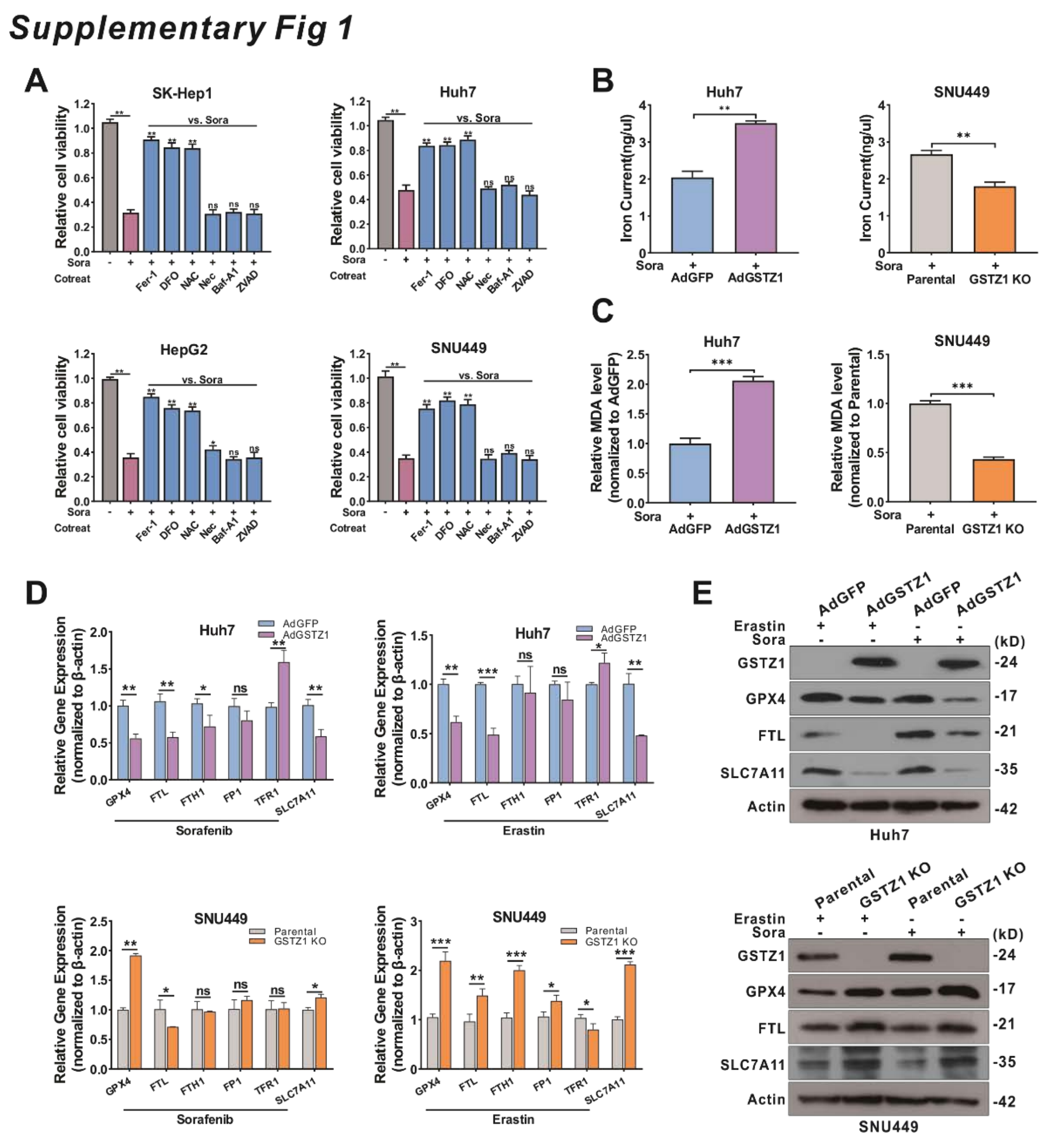


**Supplementary Figure 1.** GSTZ1 overexpression enhances sorafenib-induced ferroptosis in HCC. **a** Cell viability in HCC cells treated with sorafenib alone or in combination with Fer-1, DFO, NAC, Nec, Baf-A1, ZVAD. **b-c** The intracellular iron (**b**) and MDA (**c**) levels in GSTZ1-OE and GSTZ1-KO cells treated with sorafenib for 24 h. **d-e** mRNA (**d**) and protein (**e**) levels of target genes associated with ferroptosis in GSTZ1-OE and GSTZ1-KO cells treated with sorafenib or erastin, determined via qRT-PCR. For Western blotting, 50 μg protein was loaded per well. HCC: hepatocellular carcinoma, DMSO: dimethyl sulphoxide, Sora: sorafenib, MDA: malondialdehyde. Fer-1: ferrostatin-1, DFO: deferoxamine, NAC: N-acetyl-L-cysteine, Nec: Necrosulfonamide, Baf-A1: Bafilomycin A1, ZVAD: ZVAD-FMK. Values represent the mean ± SD (n = 3, performed in triplicate). ns: no significant difference, $*p < 0.05$, $**p < 0.01$, $***p < 0.001$, Student's t-test (two groups) or one-way ANOVA followed by Tukey tests (three groups).

### *Supplementary Fig 2*

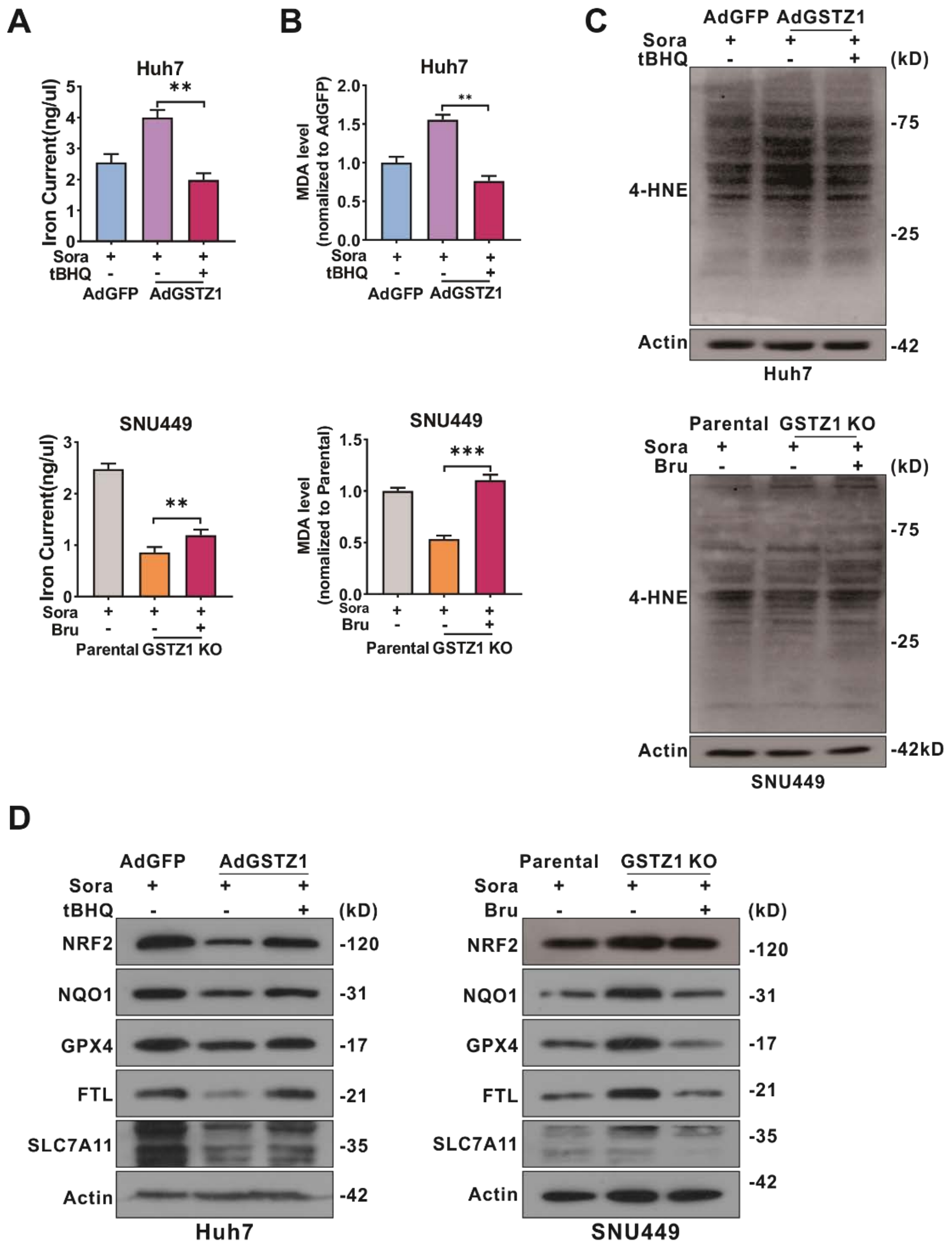


**Supplementary Figure 2.** GSTZ1 sensitizes hepatoma cells to sorafenib-induced ferroptosis through the NRF2 signaling pathway. **a-b** Levels of iron (**a**), and MDA (**b**) in GSTZ1-OE cells treated with sorafenib alone or in combination with tBHQ (top) and GSTZ1-KO cells treated with sorafenib alone or in combination with Bru (bottom). c-d The levels of 4-HNE modification and protein related with ferroptosis in GSTZ1-OE and -KO cells. The cell processing is described as above. For Western blotting, 50 μg protein was loaded per well. tBHQ: tertiary butylhydroquinone, Bru: brusatol, Sora: sorafenib, MDA: malondialdehyde, 4-HNE: 4-hydroxy-2-nonenal. Values represent the mean ± SD (n = 3, performed in triplicate). *$p < 0.05$, **$p < 0.01$, ***$p < 0.001$, Student's t-test (two groups) or one-way ANOVA followed by Tukey tests (three groups).

### *Supplementary Fig 3*

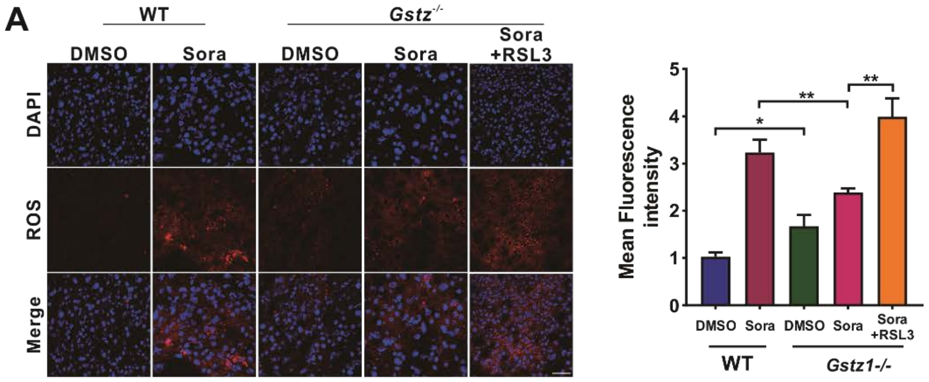


**Supplementary Figure 3.** RSL3 enhances the anticancer activity of sorafenib in Gstz1-knockout mice. **a** Representative fluorescence staining of ROS with CellROX Orange probe in hepatic tumors of five groups mice (left). Intracellular ROS quantification (right). Values represent the mean ± SD (n = 3, performed in triplicate), *p < 0.05, **p < 0.01, one-way ANOVA followed by

Tukey tests (five groups).

**Table S1 Quantitative RT-PCR Primer Sequences**

| Name | Accession Number | Source | Sequence (5'-3') |
|---|---|---|---|
| ***GSTZ1* (Human)** | NM_145870 | TsingKe Biological Technology | Forward: CCTGAAGCAAGTGGGAGAGG<br>Reverse: TGATGGTAGGGTAGGGGGTG |
| ***FTL* (Human)** | NM_000146.4 | TsingKe Biological Technology | Forward: GATGATGTGGCTCTGGAAGGC<br>Reverse: TGTGGAGGTTGGTCAGGTGG |
| ***GPX4* (Human)** | NM_002085.5 | TsingKe Biological Technology | Forward: CCGCCTTTGCCGCCTAC<br>Reverse: TTTACTTCGGTCTTGCCTCACT |
| ***FTH1* (Human)** | NM_002032.3 | TsingKe Biological Technology | Forward: CGCCAGAACTACCACCAGG<br>Reverse: CAAAGAAGTCCTCCAGCTTG |
| ***FP1* (Human)** | NM_014585.6 | TsingKe Biological Technology | Forward: TCATCGGCTGTGGCTTTATT<br>Reverse: CTGGGAGGCACAAGTAGGCT |
| ***TFR1* (Human)** | NM_003234.4 | TsingKe Biological Technology | Forward: GCTTTCCCTTTCCTTGCATAT<br>Reverse: CACGAACTGACCAGCGACCT |
| ***SLC7A11* (Human)** | NM_014331.4 | TsingKe Biological Technology | Forward: TTTCTGAGCGGCTACTGGG<br>Reverse: CAAAGGGTGCAAAACAATAACA |
| ***β-actin* (Human)** | NM_001101 | TsingKe Biological Technology | Forward: AGGCCAACCGCGAGAAGATGACC<br>Reverse: GAAGTCCAGGGCGACGTAGCAC |
| ***Gpx4* (Mouse)** | NM_008162.4 | TsingKe Biological Technology | Forward: GCAATGAGGCAAAACTGACG<br>Reverse: CCCTTGGGCTGGACTTTCA |
| ***β-actin* (Mouse)** | NM_007393 | TsingKe Biological Technology | Forward: CGTTCAATACCCCAGCCATG<br>Reverse: GACCCCGTCACCAGAGTCC |

**Supplementary Table 1.** Primer sequences used in this study.
